# From autopoiesis to self-optimization: Toward an enactive model of biological regulation

**DOI:** 10.1101/2023.02.05.527213

**Authors:** Tom Froese, Natalya Weber, Ivan Shpurov, Takashi Ikegami

## Abstract

The theory of autopoiesis has been influential in many areas of theoretical biology, especially in the fields of artificial life and origins of life. However, it has not managed to productively connect with mainstream biology, partly for theoretical reasons, but arguably mainly because deriving specific working hypotheses has been challenging. The theory has recently undergone significant conceptual development in the enactive approach to life and mind. Hidden complexity in the original conception of autopoiesis has been explicated in the service of other operationalizable concepts related to self-individuation: precariousness, adaptivity, and agency. Here we advance these developments by highlighting the interplay of these concepts with considerations from thermodynamics: reversibility, irreversibility, and path-dependence. We interpret this interplay in terms of the self-optimization model, and present modeling results that illustrate how these minimal conditions enable a system to re-organize itself such that it tends toward coordinated constraint satisfaction at the system level. Although the model is still very abstract, these results point in a direction where the enactive approach could productively connect with cell biology.

## 1 Introduction

What is life? Substantial scientific progress has been made in answering this question, especially in terms of a detailed deconstruction of the living cell into all of its whirring parts. However, when viewed from recent advances of the enactive approach, a more fundamental conception of the phenomenon of life itself is only beginning to be developed (Di Paolo, 2018). Given that for many researchers the science of the phenomenon of life is a success story – case closed – it is refreshing to hear a sobering assessment by scientists working at the frontier of biochemistry. Lane opens his book on the citric acid or Krebs cycle with a retrospective on the development of his field since the introduction of microscopes in the 17^th^ century:

“we now know what most of these whirring parts do, what they’re made of, how they function. We have taken them apart, in centrifuges or with optical tweezers, read out the code that specifies their structures, deciphered the regulatory loops that lend an illusion of purpose, listed all their parts. And yet underneath it all, we are barely any closer to understanding what breathes life into these flicks of matter. How did they first emerge from the sterile inorganic Earth? What forces coordinate their exquisite behavior? Do they experience any sort of feelings?” (Lane, 2022, pp. 3-4)

Without denying the importance of genes, Lane puts his finger on the regulation of energy flows as the most fundamental mark of the living, which enables a cell to continuously regenerate itself biochemically from simpler components. Akin to genetic heredity, this ongoing metabolic process has been passed from generation to generation, thereby linking us all the way back to the origins of life. At the core of this process is a cycle of transformation of energy and matter – the Krebs or citric-acid cycle – whose molecular basis is by now well described but whose deeper meaning remains elusive. According to Lane, this cycle of energy and matter “conceals a strained balance of opposites, a yin and yang, which touches on all aspects of life” (ibid. p. 19). This is because the Krebs cycle is both the engine of biosynthesis, driving cellular growth and renewal, while also being capable of – though never at the same time -burning those same molecules in respiration, driving energy generation instead. This arrangement means that there is a coordination problem that requires an adaptive solution:

“The problem in all these disparate cases is how to balance the yin and yang of the Krebs cycle – how to offset the needs of energy generation against the synthesis of new organic molecules. This question brings the Krebs cycle into sharp modern focus but does not explain why the cycle has a yin and yang at all.” (Lane, 2022, p. 21)

In other words, why is there this apparently unnecessary tension? Is it just an evolutionary epiphenomenon that there happens to be a metabolic cycle that, depending on the direction of its operation, can service one of two requirements, i.e., either component production or energy generation? Or does this paradoxical organization, which entails that the cycle can never meet these two requirements simultaneously, but must coordinate them over time, reflect a deeper dialectic at work in life? Lane’s extensive investigation into these issues leads him to propose that a cycle of yin and yang pays off in terms of efficiency and flexibility, for example. In addition, his intuition is that there is a link between this paradoxical organization and the qualitative side of regulation, namely striving, feeling, self, and even consciousness. Nevertheless, he stops short of providing a theoretical framework that could conceptually integrate this life-mind continuity.

The possibility that the organization of this metabolic pathway into a cycle of yin and yang is an essential part of its function, rather than merely a frozen evolutionary accident of some sort, is in line with conceptual advances in a distant field – the enactive approach, which is developing a new biological foundation for cognitive science by drawing inspiration from autopoietic theory, systems biology, and phenomenological philosophy (Thompson, 2007). In the following we first advance this biological foundation by deriving a potential mechanism of unsupervised biological regulation – self-optimization – from its abstract conception of life. We then demonstrate that this mechanism has a workable model by implementing a mathematical computer simulation, which enables us to explore the minimal conditions of adaptive constraint satisfaction. Although these contributions are mainly conceptual, they help to make intelligible how life could benefit from an itinerancy between two opposing requirements: rather than posing a problem of balance, this irreducible tension is what drives adaptive coordination in the first place.

## 2 From autopoietic theory to the enactive approach

The autopoietic tradition is well known for attempting to provide an operational definition that categorically separates living from non-living systems in terms of their self-production as a physically bounded individual (Varela, Maturana, & Uribe, 1974). An intuitive sense of the key idea is provided by contrasting allopoietic and autopoietic systems, for example a factory that produces something other than itself compared to a cell which continually regenerates itself. Subsequent developments of the autopoietic tradition derive a continuity from life to mind to sociality (Maturana & Varela, 1987; Varela, 1979). Although we do not concern ourselves with these extensions here, they serve to highlight that a revision or clarification of autopoiesis can have repercussions for a broad range of phenomena. A canonical definition of autopoiesis that has become influential in recent developments of enactive cognitive science reads as follows:

> “An autopoietic system is organized (defined as a unity) as a network of processes of production (synthesis and destruction) of components such that these components:
>
> (i)continuously regenerate and realize the network that produces them, and
>
> (ii)constitute the system as a distinguishable unity in the domain in which they exist.” (Varela, 1997, p. 75)

Originally, autopoietic theory was a purely mechanistic theory that adopted an anti-teleological stance and kept the domain of the living system’s physiology strictly distinct from the domain of behavior (Maturana & Varela, 1980). However, it underwent substantial transformations in the context of the enactive approach: autopoiesis under precarious conditions was taken to be the biological basis for intentionality, giving rise to an intrinsic goal-directedness that is accompanied by a perspective of concern – primarily for the continuation of the process of self-production (A. Weber & Varela, 2002). Yet the all-or-nothing categorical concept of autopoiesis is not sufficient to explain how a living system can exhibit sense-making that depends on a graded sensitivity with respect to how well it satisfies its intrinsic goals (Di Paolo, 2005). This capacity for sense-making also requires a living system to be adaptive, such that it can make corrective changes to its internal or behavioral dynamics with respect to its boundary of viability before that boundary is crossed – because otherwise there would no longer be an individual left to respond adaptively.

Subsequent theoretical developments of the enactive approach showed how precariousness and adaptivity are no mere contingent add-ons to autopoiesis, but can be understood as essential aspects of life’s concrete embodiment (Di Paolo, 2018). The physical basis of autopoiesis under thermodynamically far-from-equilibrium conditions is characterized by an irreducible tension at the core of living systems between two opposing physical requirements, in which their boundary in the form of a semipermeable membrane plays an essential role:

1. *Closure:* The requirement of a living system to maintain its systemic identity as an autonomous individual that is physically distinct from its environment (e.g., by closing its boundary to certain kinds of material and energetic exchanges with its environment), and
2. *Openness:* The requirement of a living system to obtain free energy from its environment to work on maintaining its network of processes (e.g., by opening its boundary to certain exchanges of matter and energy with its environment).

To maximally satisfy these two requirements simultaneously is in principle impossible. The living system is therefore always torn between two mutually exclusive requirements with respect to how it relates with its environment, namely, to paradoxically strive for total closure and total openness (Di Paolo, Buhrmann, & Barandiaran, 2017). If a living system tends too much in either direction it will cease to live: it will starve because of unmet energy needs, or dissolve into thermodynamic equilibrium with its environment, respectively. This entails that there is no stable relational state in which a living system can simply abide. As such, precariousness is inherent in living systems: it is ultimately based on the impossibility of *simultaneously* satisfying the two physical requirements of maximizing closure and openness with respect to its environment. Accordingly, any living system has to have the capacity to survive its own inherent precariousness by flexibly coordinating the satisfaction of these divergent requirements over time. It is therefore tempting to attribute to such minimal living systems not only the capacity of adaptivity, but perhaps even of agency (see Figure 1).

**Figure 1.**
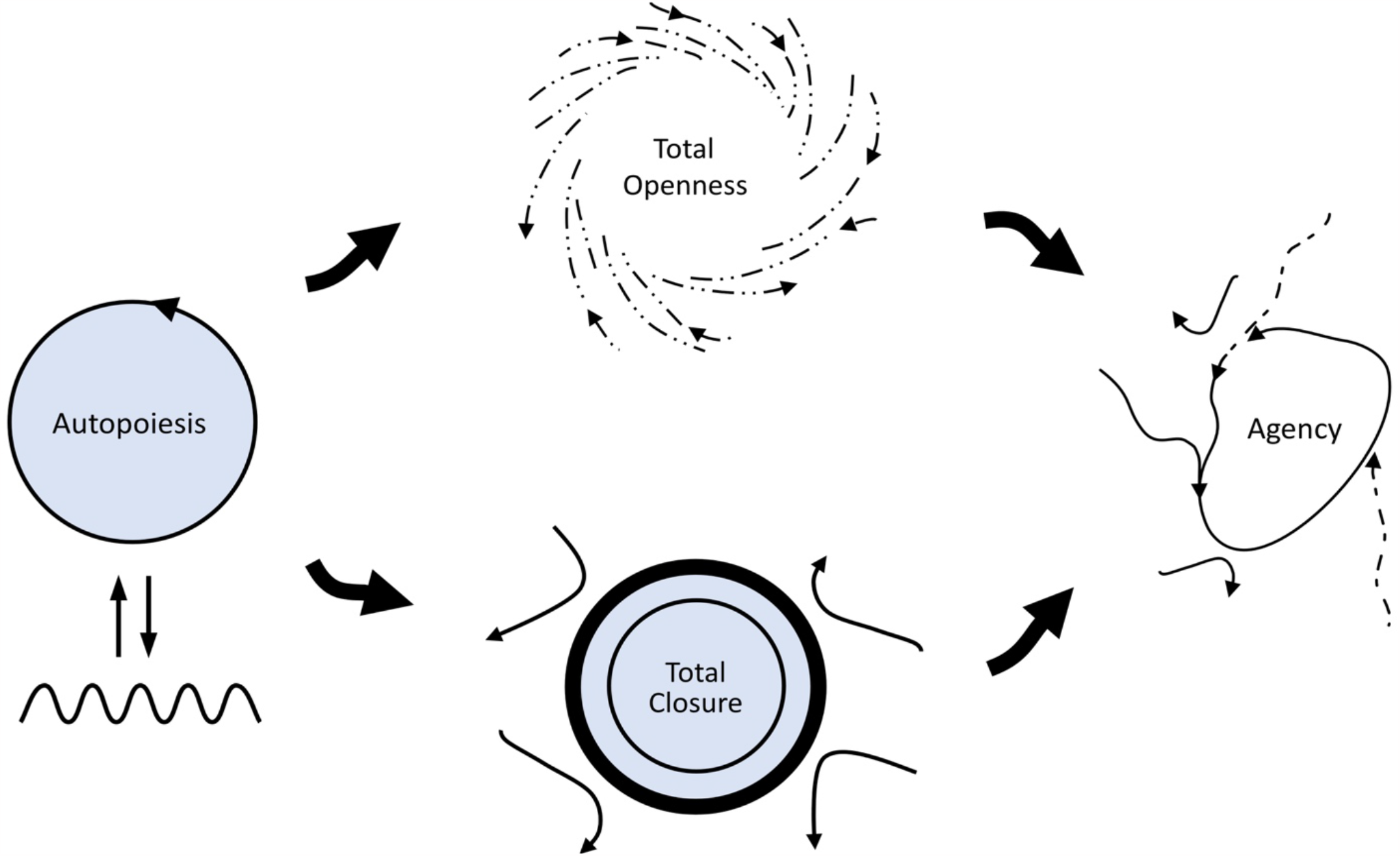
The primordial tension of self-individuation according to the enactive conception of life. On the left, there is the classical Maturana and Varela diagram of autopoiesis: a distinct self-producing individual (circular arrow) in structural coupling (straight arrows) with its environment (wavy line). In the middle, the two opposing requirements are depicted: on the top, the ideal realization of energetic flows required for self-production tends toward total openness; on the bottom, the ideal realization of the boundary required for self-distinction tends toward total closure. On the right, the resolution of this tension is depicted in terms of an agent’s adaptive regulation: a partial satisfaction of the two requirements is coordinated over time. Redrawn from Di Paolo, Buhrmann, and Barandiaran (2017, p. 135).

However, for the minimal kind of living system that we are considering here, an attribution of adaptivity or even agency is in danger of setting the bar rather high, especially if we assume that the capacity involves some kind of top-down supervision or control mechanism. Consider, for example, how adaptivity has been canonically defined in the enactive approach:

> “Robustness implies endurance but not necessarily *adaptivity* which is a special manner of being tolerant to challenges by actively monitoring perturbations and compensating for their tendencies. Adaptivity is defined as:
>
> A system’s capacity, in some circumstances, to regulate its states and its relation to the environment with the result that, if the states are sufficiently close to the boundary of viability,
>
> 1. Tendencies are distinguished and acted upon depending on whether the states will approach or recede from the boundary and, as a consequence,
> 2. Tendencies of the first kind are moved closer to or transformed into tendencies of the second and so future states are prevented from reaching the boundary with an outward velocity.”
>
> (Di Paolo, 2005, p. 438; Di Paolo et al., 2017, p. 122)

Indeed, a self-producing system that can resist succumbing to the irresolvable tension between openness and closedness by coordinating these conflicting tendencies over time requires more than mere robustness. But must it require a process of actively monitoring based on capacities for distinguishing and regulating state trajectories with respect to its boundary of viability? To be fair, there is the caveat that “this capacity may result from the action of dedicated mechanisms or it may be an emergent aspect of specific ways of realizing autopoiesis” (Di Paolo, 2005, p. 438). However, if we opt for the latter “emergent” interpretation and apply it to the primordial tension of self-individuation, we basically turn this definition of adaptivity into a re-description of the living system’s temporal dynamics: whenever the system comes too close to one of its mutually exclusive requirements – too much openness or too much closedness – it tends to turn back in the other direction. But how does the system manage to turn back? It seems that a living agent requires a dedicated regulatory mechanism that is at least partially decoupled from its metabolic self-production (Barandiaran & Moreno, 2008).

One possible mechanism inspired by cybernetics is ultrastability (Ashby, 1960): the basic idea is that when one of the system’s essential variables exceeds its boundary of viability, it triggers a stochastic search through its parameter space until it comes across a parameter that returns it to a stable state. Notably, ultrastability likely served as an important inspiration for the original work on autopoiesis, namely by taking the living system’s own organization as the essential variable to be maintained (Froese & Stewart, 2010). However, this brings us back to the same problem: this generalization of ultrastability to life itself would entail that the system needs to cross its boundary of viability for the process of adaptation to begin. In other words, in this case there could be no possibility of directed lifetime adaptation, which would indeed be consistent with the traditional autopoietic theory’s notion of natural drift (Maturana & Mpodozis, 2000).

In contrast, Varela (1992) later explicitly conceived of autopoiesis as the starting point of a biology of intentionality, that is, precisely of directedness. For this purpose, the concept of ultrastability may still be acceptable for biological variables that are not too essential after all, such as ranges of neural firing rates (Di Paolo, 2003), but it does not work for strictly essential variables whose boundary of viability coincides with the very distinction between life and death. Consider body temperature as a typical essential variable: a mechanism that would require an organism to die from overheating before it could start a stochastic search for a way of cooling down would be of little use as a model of adaptive regulation.

It may seem that ultrastability has a straightforward solution: set the threshold for starting the stochastic search to be within the boundary of viability. However, how could a minimal living system, which consists of not much more than autopoiesis, adaptively regulate its operations in relation to its own boundary of viability, given that it must accomplish this without the benefit of complex mechanisms fine-tuned by long-term evolutionary optimization? More precisely, how could it work out what would be its optimal state given current conditions and then work out what should be its next state given its current state? Again, the use of error or reward signals set the bar rather high for the kind of minimal living system we are considering here. What is needed to get adaptivity, and hence life itself, started is a basic mechanism for coordinated constraint satisfaction, ideally capable of implicit generalization or anticipation, but which does not depend on active monitoring and comparing. Finding such a basic mechanism is conceptually challenging; most artificial optimization mechanisms depend on assessments of how “good” the current state is, e.g., backpropagation error, prediction error, reward, etc. They are therefore not applicable to minimal autopoietic systems to which we cannot attribute such knowledge.

## 3 Adaptivity as self-optimization

Fortunately, there exists a mathematical model of adaptive change that is unsupervised and distributed, and that does not rely on any reward nor error function, and which hence could fit the bill. This model was first introduced by Watson and colleagues under the label of “self-modeling” (Watson, Buckley, & Mills, 2011; Watson, Mills, & Buckley, 2011a, 2011b), and continues to find a wide range of theoretical applications (Watson, Levin, & Buckley, 2022; Watson & Szathmáry, 2016). The model was later renamed as the “Self-Optimization” (SO) model by Froese and colleagues (Zarco & Froese, 2018) to avoid any unintended associations with the traditional cognitive science concept of an internal representational self or world model, which would set the bar too high again. We continue this alternative naming convention here, and return to conceptual considerations of its status as a model in the discussion section.

The SO model is best understood by considering a classic Hopfield (1982) neural network. Each node of the network has a binary state whose value is updated in accordance with the states of the other nodes to which it is connected, and by taking into account the constraints that are placed on their relationships. These constraints between nodes are specified in terms of the weights of the network’s connections. A complex weight matrix specifies the contours of a state space, which can be imagined in appearance as similar to a hilly energy landscape. In terms of this metaphor, like a ball rolling down a slope and coming to rest as soon as it reaches the bottom, the network’s nodes change state according to their connections until the network converges to an equilibrium point – an attractor – which is equivalent to a coordination of all the states with respect to the network’s constraints. The overall success of this coordination can be measured in terms of a suitable energy function, according to which lower energy indicates better constraint satisfaction. However, this distributed coordination process is not sufficient to deal with a complex constraint satisfaction problem: it is overwhelmingly likely that the network will come to rest in a local equilibrium, that is, in a state configuration that only poorly satisfies constraints at the level of the whole. Even if the network is repeatedly reset to a different initial state and allowed to reconverge, optimal solutions to complex constraint problems are unlikely to be found in reasonable time.

The key insight by Watson and colleagues was that this limitation can be addressed effectively by adding a basic form of *plasticity*: associative learning. Specifically, they used a distributed, unsupervised form of correlational learning that in the field of neuroscience was popularized as Hebb’s (1949) rule, namely that “neurons that fire together wire together”, and which was successfully modeled in Hopfield artificial neural networks. In this context, Hebbian learning works akin to a principle of precedent: it involves making changes to the weights of the network such that past state correlations are more likely to reoccur in future states. When an attractor is visited, its basin of attraction is enlarged, making it more likely for that attractor to be visited again in the future. Importantly, if there is some structural regularity to exploit, the basins of other attractors with overlapping features also get enlarged according to the amount of overlap, thereby giving rise to a form of associative memory. In other words, a Hopfield network that has the capacity to repeatedly escape from local equilibria, and which slowly adjusts its connections according to Hebbian learning, has a range of useful properties: for example, it is robust to noise, and it can generalize over past states in such a way that it becomes overwhelmingly likely to converge on deeper equilibrium state configurations that would otherwise be practically inaccessible (Watson, Buckley, et al., 2011; Watson, Mills, et al., 2011b).

This SO model is particularly attractive as a possible mechanism at work in minimal living systems, since it has only very basic requirements that can be met by various other artificial neural network architectures (e.g., Woodward, Froese, & Ikegami, 2015; Zarco & Froese, 2018), and also by other complex networks that are not typically thought of in cognitive terms, such as gene regulation networks (e.g., Watson, Wagner, Pavlicev, Weinreich, & Mills, 2014) and social networks (e.g., Davies, Watson, Mills, Buckley, & Noble, 2011; Froese, Gershenson, & Manzanilla, 2014; Froese & Manzanilla, 2018). There are different ways of ensuring that the systems can escape from their local equilibria, some of which will be domain specific. For example, in the case of gene regulation networks and ecological networks it could be due to environmental fluctuations (Watson et al., 2022), in the case of social networks it could also be anti-structure temporally enacted in community ritual (Froese, 2018), while neural networks could be benefitting from avalanche dynamics that are associated with self-organized criticality (Shpurov & Froese, 2021). Note also that associative memory is a general principle that does not depend on neural plasticity; associative memory can also be implemented without changes to the network’s connections (Izquierdo, Harvey, & Beer, 2008). It could in principle be based on a very minimal mechanism; for example, associative learning of temporal correlations is even exhibited by basic reaction-diffusion chemistry (Bartlett & Louapre, 2022).

In summary, viewed from a higher level of abstraction, the SO model depends on three kinds of processes that would need to be satisfied by a biological mechanism (see also Figure 2):

1. *Closed dynamics*: maintenance of a specific meta-stable state that is characterized by increased coordination among the constraints of the system, and
2. *Open dynamics*: spontaneous transient through state space, in a way that increases chances of convergence into a different meta-stable state, and
3. *Plasticity*: sedimentation of correlations of previously visited meta-stable states, in a way that increases the chances of those correlations to reoccur.

**Figure 2.**
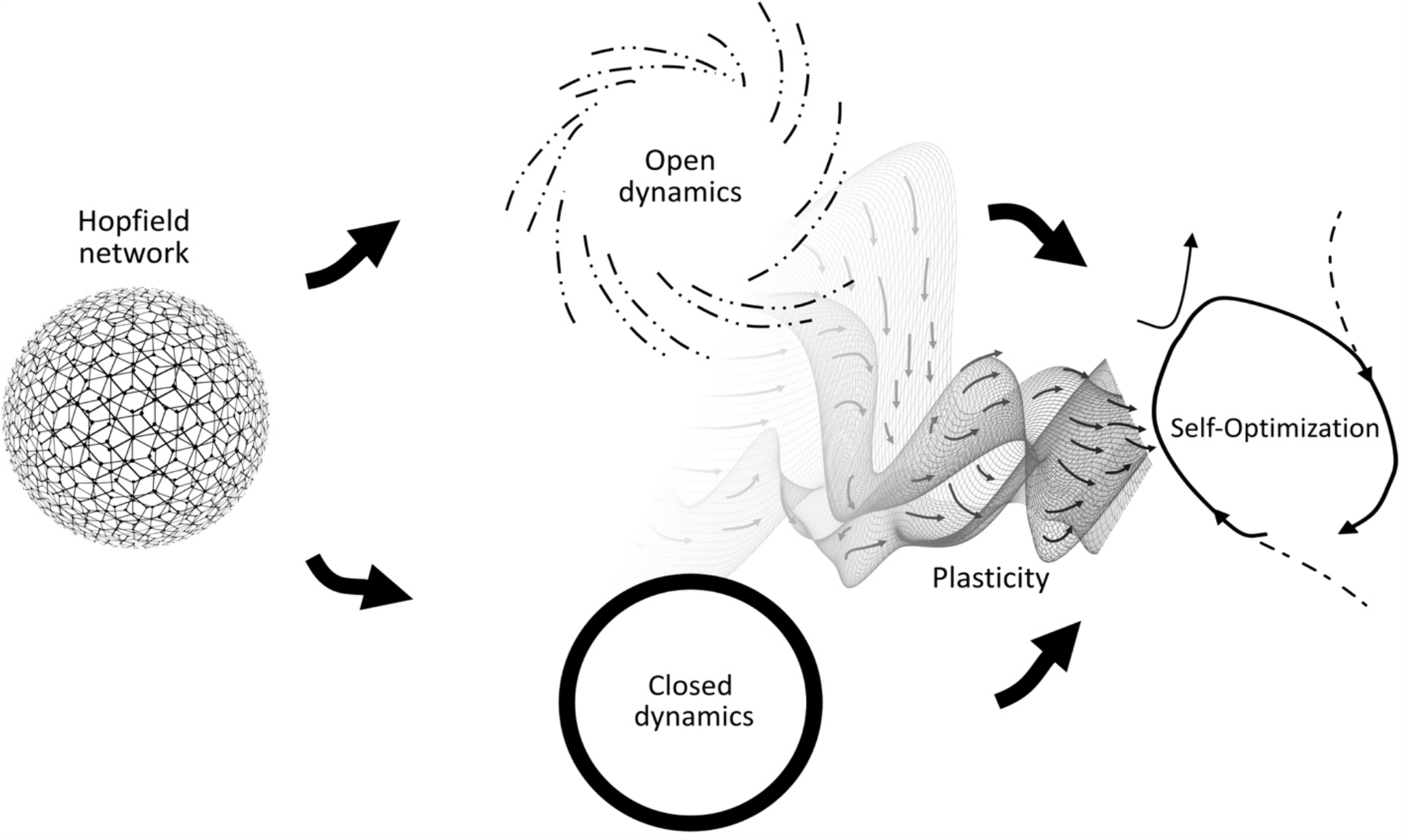
Schematic of the Self-Optimization (SO) model. On the left, there is the classical Hopfield network. The weights of the connections between nodes can represent the elements of a constraint satisfaction problem. The states tend to stabilize in an ordered configuration, which at the same time represents a possible solution to that problem, albeit for complex problems this is typically only locally optimal. In the middle, the two modes of the SO model are depicted: on the top, constraints on the network state are bracketed and it enters a transient; on the bottom, constraints force the network state to stabilize in another coordinated configuration. On the right, the addition of plasticity to the network’s itinerancy between these two modes, implemented in the form of unsupervised associative learning, gives rise to self-optimization: over time, the network re-organizes such that it becomes more likely to select paths toward greater coordination from the set of visited configurations. It can even generalize over these past configurations such that it begins to select rare paths toward the most highly coordinated configurations that would otherwise remain unvisited in practice.

All three aspects of the SO model are already implicit in the enactive conception of life, with its emphasis on the living system’s precarious itinerancy between the two tendencies toward total openness and total closure in a path-dependent manner. This link can be made more explicit by introducing thermodynamic considerations, and a particularly suitable concept for this purpose is dissipative adaptation in driven self-assembly (England, 2015). When the living system drives itself toward total openness, this is equivalent to increasing its energy, while closing its boundary has the opposite effect, namely decreasing its energy. This actively regulated change in the living system’s level of energy is akin to a thermodynamically open system that is rhythmically driven through a specific sequence of configurations by a time-varying field. Effectively, this is a form of driven barrier hopping: a high point in energy momentarily lowers the activation energy required to overcome energy barriers standing in the way of transitioning into another configuration, hence increasing the likelihood that a barrier is crossed during those moments. A transition involves dissipation of heat into the environment, which has the consequence of decreasing the likelihood that the system will return to its previous configuration. Accordingly, while the energy rhythmically reverses back and forth between high and low values, the specific sequence of transitions between configurations is path-dependent and statistically irreversible. In addition, there is a form of plasticity: the more energy is absorbed from the environment and then dissipated during transition to another configuration, the more irreversible is that transition.

“Thus, while any given change in shape for the system is mostly random, the most durable and irreversible of these shifts in configuration occur when the system happens to be momentarily better at absorbing and dissipating work. With the passage of time, the ‘memory’ of these less erasable changes accumulates preferentially, and the system increasingly adopts shapes that resemble those in its history where dissipation occurred. Looking backward at the likely history of a product of this non-equilibrium process, the structure will appear to us like it has self-organized into a state that is ‘well adapted’ to the environmental conditions. This is the phenomenon of dissipative adaptation.” (England, 2015, p. 922)

According to the enactive approach, therefore, a minimal living system has all three essential ingredients needed for the spontaneous self-optimization of its organization – closed dynamics, open dynamics, and plasticity – such that its states will be locally coordinated in a way that ends up satisfying more constraints at the system level. Plasticity in particular plays a key role in the self-organization of adaptivity based on these itinerant dynamics, both in the SO model and in the phenomenon of dissipative adaptation. Accordingly, we propose to make its importance for the enactive approach to autopoiesis more explicit by graphically updating the conceptual diagram originally proposed by Di Paolo and colleagues (Figure 3). This change is in line with a growing emphasis on historicity as a distinguishing element of the enactive conception of life (Di Paolo, Thompson, & Beer, 2021).

**Figure 3.**
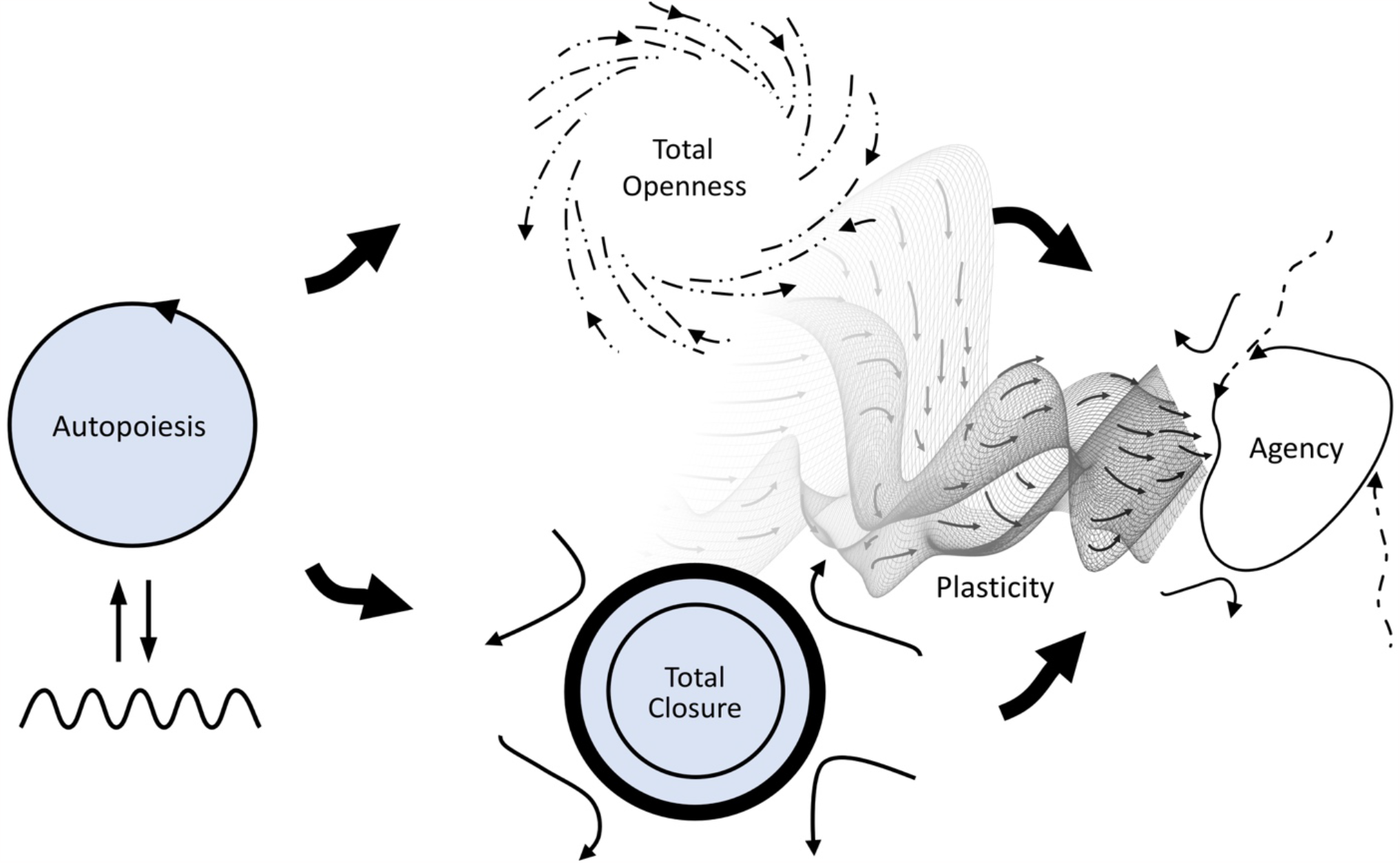
Updated schematic of the primordial tension of self-individuation according to the enactive conception of life. See the caption of Figure 1 for details. On the right, it is now made explicit that the resolution of this tension requires a historical dimension: an agent’s adaptive regulation of the itinerancy between satisfying these two requirements additionally depends on the accumulation of plastic changes during that itinerancy that make some state configurations more likely. Redrawn and revised from Di Paolo, Buhrmann, and Barandiaran (2017, p. 135).

In the next two sections we will illustrate this self-organization of adaptivity at an abstract level of description with a new implementation of the SO model. We investigated two open questions:

1. *Emergent process versus dedicated sub-system*: Di Paolo (2005) allowed that adaptivity was either an emergent aspect or the result of a dedicated mechanism, but subsequent work argued for the need of a regulatory mechanism that is partially decoupled from self-production (Barandiaran & Moreno, 2008). It has also been proposed that the SO model can work in two modes: i) by accumulating changes directly at the system level such that the SO emerges out of the whole, ii) or by capturing changes in a dedicated sub-system such that it can later function in a stand-alone manner (Watson, Buckley, & Mills, 2009). Here we systematically tested this proposal for a dedicated subsystem.
2. *Complete versus partial resets*: Given our association of the tendency to openness with the living system driving itself to a new configuration, the traditional implementation of this transition in the SO model as a complete instantaneous re-initialization to an arbitrary state configuration is problematic. Support for the feasibility of using only partial and temporally extended resets comes from a SO model with local resets that are dependent on neural avalanche dynamics (Shpurov & Froese, 2021). Here we systematically tested the effects of the amount and duration of resets. It is not clear how to precisely model the shape of the transition; we considered two general possibilities: (a) that the system switches immediately (rectangular pulse), and (b) that the system switches gradually (sinusoidal pulse).

A word of caution before we start discussing the details of the SO model: As highlighted above, even though we propose that is fruitful to interpret the SO model in thermodynamic terms as a form of dissipative adaptation, this interpretation remains highly abstract. The SO model is not implemented in a spatially embodied manner that would allow for consideration of the role of an environmental boundary that is capable of regulating energy gradients. Future work could take inspiration from artificial life models that link homeostatic temperature regulation to the emergence of sensorimotor loops in mobile agents (Ikegami & Suzuki, 2008). More generally, it is an outstanding challenge to demonstrate that the SO model can be implemented in terms of a coupled agent-based model to regulate behavior (Zarco & Froese, 2018).

## 4 The Self-Optimization (SO) model

In the original SO model (Watson, Buckley, et al., 2011), the system is a Hopfield network with *N* nodes, where each node can be in one of two possible states, either *s* = 1 or *s* = −1. A vector ***S***(*t*) = {*s*_1_(*t*), …, *s*_*N*_(*t*)} defines the state of the system at any given time. Overall, there are 2^*N*^ possible states of the system. Initial weight matrix **W**_0_ of size *N* × *N* defines the connections between the nodes. The initial weight matrix **W**_0_ is chosen to represent the constraints of a constraint optimization problem (see Sec. 4.1), and the connections of the weight matrix **W**_0_ are chosen to be symmetric. The system is updated asynchronously, that is a random node is selected at each time step and updated according to the following rule

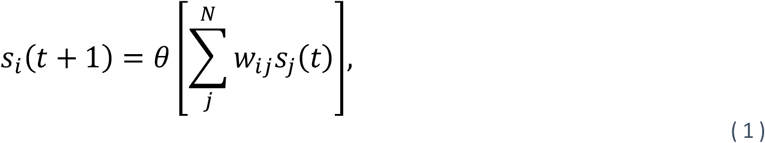

***W***where *w*_*ij*_ are elements of the weight matrix **W** (Eq. (*2*)), and *θ* is a Heaviside threshold function (taking values -1 for negative arguments and +1 otherwise). During the learning stage, the weights are updated according to a Hebbian learning rule

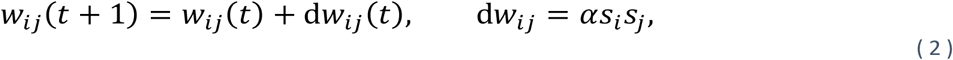

where *α >* 0 is a learning rate constant. In a vector form Eq. (2) can rewritten as

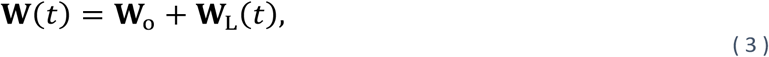

where **W**_L_ is the accumulation of Hebbian learning updates onto the original weights. Thus without learning **W**(*t*) = **W**_0_. The energy function is computed with respect to the original weight matrix **W**_0_, and thus provides a read-out of how well the system is satisfying its initial constraints. It is defined as

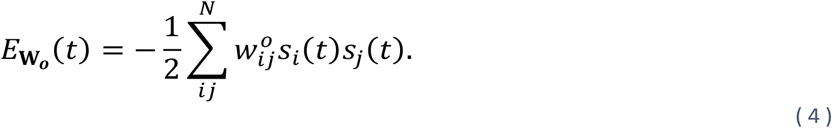

After performing *T* steps of updating the states, during which the system will converge towards some local minimum, the entire state is reset instantaneously to a new random state. The system is repeatedly allowed to converge to a local attractor for *M* resets.

### 4.1 Original weight matrix

The original weight matrix **W**_0_ represents a constraint satisfaction problem (a weighted Max-SAT problem) that the system must solve by coordinating its state to optimally resolve tensions between its constituent elements. The matrix of constraints is generated by randomly sampling from [-1,1]. To add additional complexity to the problem, in some cases modular structure is introduced by increasing the contribution from the blocks of weights on the main diagonal to the overall performance score, as it is measured by the system’s energy function. In Section 5, we present simulation results for a weight matrix that has 20 modules of size 5-by-5 with intramodule weights set at random to either 1 or -1, and intermodule weights set at random to either 0.1 or -0.1. Such a sparsely connected or modular organization enables the study of the generalization abilities of the model, as the solution which optimally resolves the system’s constraints can emerge from the combination of local optima, which resolve the constraints on the level of the individual modules. For details on various weight matrices as constraint problems, see the work by Watson and colleagues (Watson et al., 2009; Watson, Buckley, et al., 2011).

### 4.2 System’s resets

Existing work periodically resets the whole system to an arbitrary initial state all at once, which does not fit very well the idea of a system that is driven to itinerate between high and low energy.

We therefore consider probabilistic reset functions that reset only a certain number of discrete states based on a probability that oscillates over time. We examine two cases: a rectangular pulse train and a sinusoidal pulse train. The difference between these two cases lies in the percentage of the system that is reset per pulse. While for a rectangular pulse train the same percentage of the system is reset during the width of the pulse, for a sinusoidal pulse train the amount of states that will be reset depends on the height of the pulse at each time step.

#### Rectangular pulse train reset function

We use a rectangular pulse train *R*_rect_(*t*) in the following form

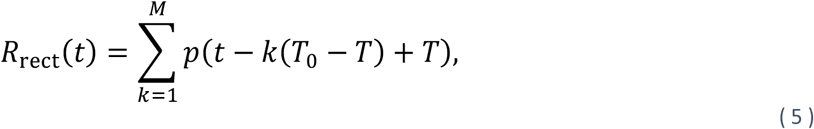

where *T*_0_ is the period, *T* is the duration or the width of a pulse, and *p*(*t*) is a rectangular pulse defined as:

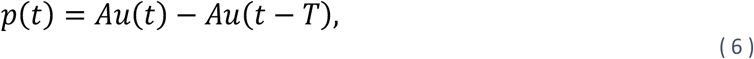

where *A* is the amplitude of the pulse, and *u*(*t*) is a unit function (taking values 0 for negative arguments and 1 otherwise).

#### Sinusoidal pulse train reset function

We define the sinusoidal pulse train *R*_*s*_(*t*) as

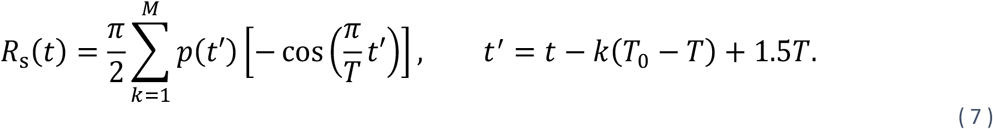

The arguments of *R*_*rect*_,(*t*) and *R*_*s*_(*t*) are chosen such that before the first pulse, after the last pulse, and between all the pulses there is the same period *T*_0_ that will allow the system to reach convergence before the next pulse. The factor *π***/**2 in Eq.(*7*) ensures that for the same values of *T*_0_, *T, A* the area under a pulse is the same for both *R*_*rect*_ (*t*) and *R*_*s*_(*t*). An example of *R*_*rect*,_(*t*) and *R*_*s*_(*t*) with 5 pulses for a system of *N* = 100 nodes is shown in Fig. 4.

**Figure 4.**
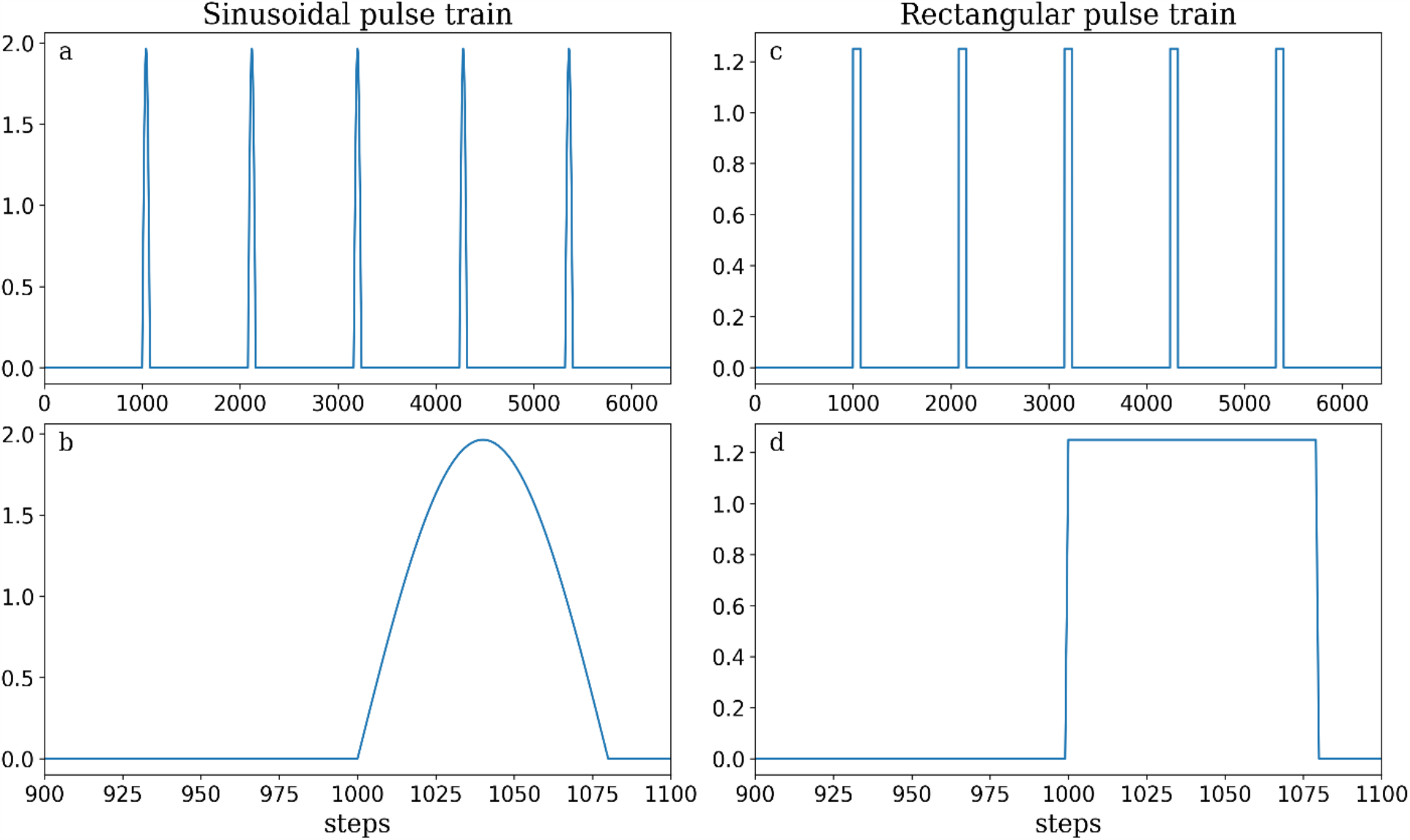
An illustration of the two functions that were selected to drive the periodic partial state resets of the Self-Optimization model. Periodic pulse trains (a,b -sinusoidal, c,d – rectangular) of 5 pulses with amplitude A = 1.25, period T_0_ = 1000, and width T = 80 for a system with N = 100 nodes. The bottom row zooms in on the first pulse to better illustrate its shape.

At each time step, the fractional part *r ∈* (0,1), and the integral part *n ∈* [0, *N*] of the height of the pulse are computed. Accordingly, *n* discrete states of the system will be reset at random to either 1 or -1, and an additional one discrete state will be reset randomly with probability *r*. For example, if the height of the pulse is 55.6, *n* = 55 discrete states *s* of the system’s state ***S*** will be reset at random to either 1 or -1, and additional one state will be reset with a probability *r* = 0.6.

## 5 Results

We first replicate a well-established finding: when a Hopfield network is repeatedly reset to an arbitrarily disordered state configuration, and from there transitions into a locally ordered state configuration where it remains for a while, Hebbian learning will induce an associative memory of the ordered states, which becomes the basis of generalized capacity for constraint satisfaction of the original problem space (section 5.1). This is an emergent capacity, yet we show that the acquired memory can also stand in for the optimized Hopfield network as a whole and does equally well in constraint satisfaction. We then go beyond the implementations explored in the existing literature, by showing that the SO model does not depend on the precise way in which the state configuration becomes disordered; different kinds of partial disordering work too (section 5.2 and Appendix A). Indeed, the results indicate that a gradual, probabilistic reset of states to arbitrary values could be preferable in terms of temporal efficiency of self-optimization, when compared to a complete instantaneous reset of all states (Appendix B).

## 5.1 Original Self-Optimization model (instantaneous reset of the entire system)

Here we present results that illustrate how the standard SO model works, i.e., with resets that instantaneously reset the system to another arbitrary initial state. In all simulations we examine a system with *N* = 100 nodes, a convergence period of 10*N* steps, a modular weight matrix with 20 modules of size 5 × 5 (where intramodule weights set at random to either 1 or -1, and intermodule weights set at random to either 0.1 or -0.1), and a learning rate *α* = 1 × 10^34^.

Figure 5 shows distributions of energy of the system at the end of a convergence and before the system is instantaneously reset to a new random state for 4000 resets: 1000 resets without learning, followed by 1000 resets with learning (Eq. 2), followed by 1000 resets without learning with the dynamics of the states driven by the final weight matrix **W**, followed by 1000 resets again without learning, but now the dynamics of the states are driven only by the learned part of the weight matrix, **W**_L_.

**Figure 5.**
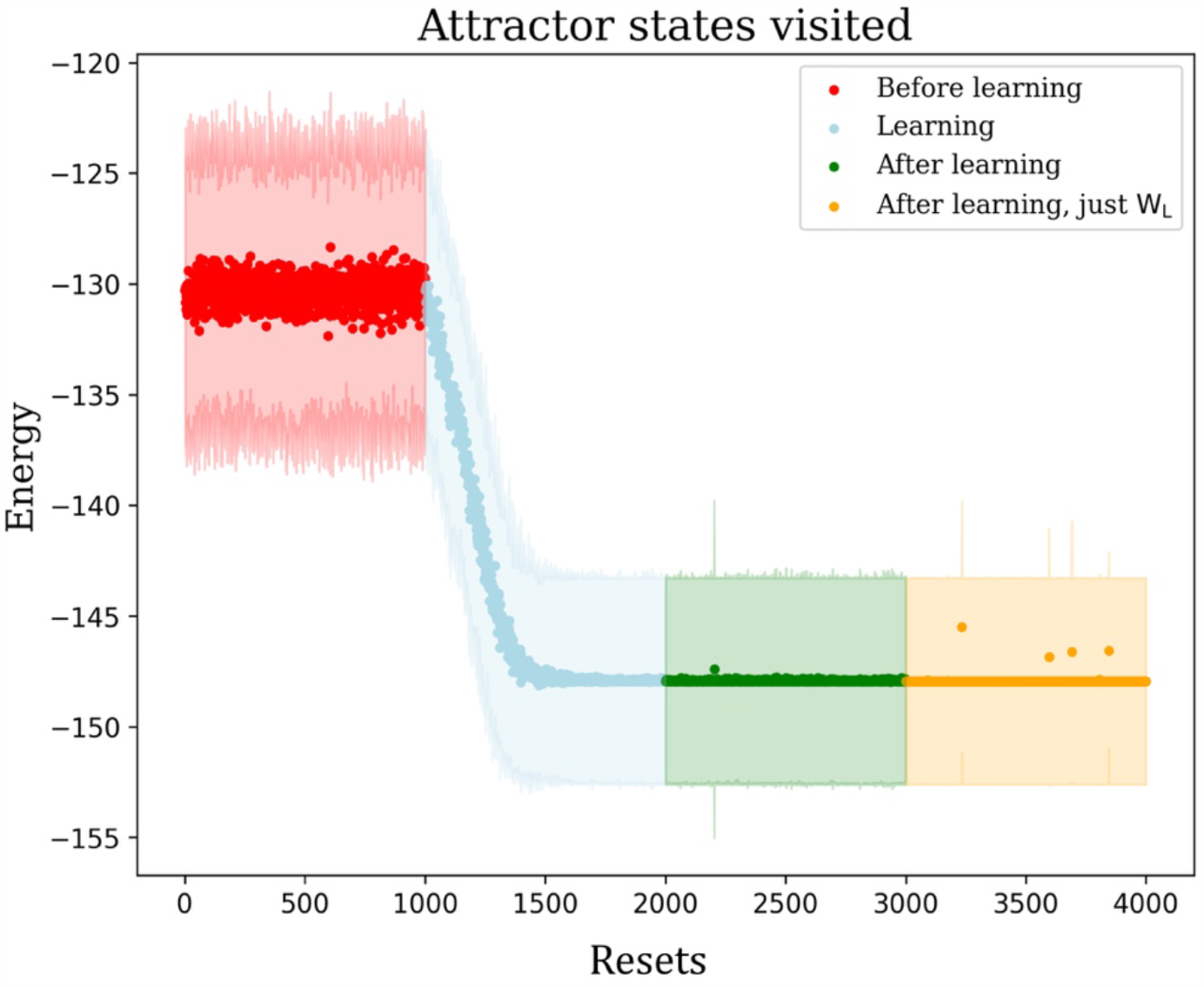
Energy of attractors visited, as computed in terms of the original constraint satisfaction problem. Each point represents the mean energy for 100 random runs at the end of convergence for a set without learning (resets 1–1000, red), during learning (1001–2000, blue), after learning with dynamics computed with the self-optimized **W** (2001-3000, green), and after learning with dynamics computed with the acquired changes **W**_L_ alone (3001-4000, orange). The shaded areas in corresponding colors denote the standard deviation of the distribution.

The first set of 1000 resets demonstrate the initial distribution of attractor states according to the randomly chosen weight matrix. The second set demonstrates the effect of learning, and the third set demonstrates how after learning the system remains in the learned configuration. We can see that during learning the system on average converges to an energy *E*_mean_∼ − 147 roughly after 520 resets. In other words, by being repeatedly perturbed out of its equilibrium state, the system is able to generalize over past equilibria and tend toward better equilibria that it would otherwise never encounter.***W***As illustrated in Figure 6, the finding shown in Figure 5 is representative of a general pattern. The standard Hopfield network has difficulties coordinating its states so that they converge on deeper equilibria (Fig. 6d), while the SO model can self-optimize such that eventually it always converges on a deep equilibrium no matter the initial state (Fig. 6e). This result has been demonstrated in a multitude of scenarios since Watson and colleagues’ (2009) original technical report. What has so far not been shown explicitly, but was discussed conceptually in that report, is that the same behavior can be observed by the learned weights **W**_L_ that “come to dominate the behaviour of the system” (ibid.), as shown by the fourth set of 1000 resets. Thus, the learned weights can stand in for the whole optimized weight matrix with comparable results (Fig. 6f).

**Figure 6.**
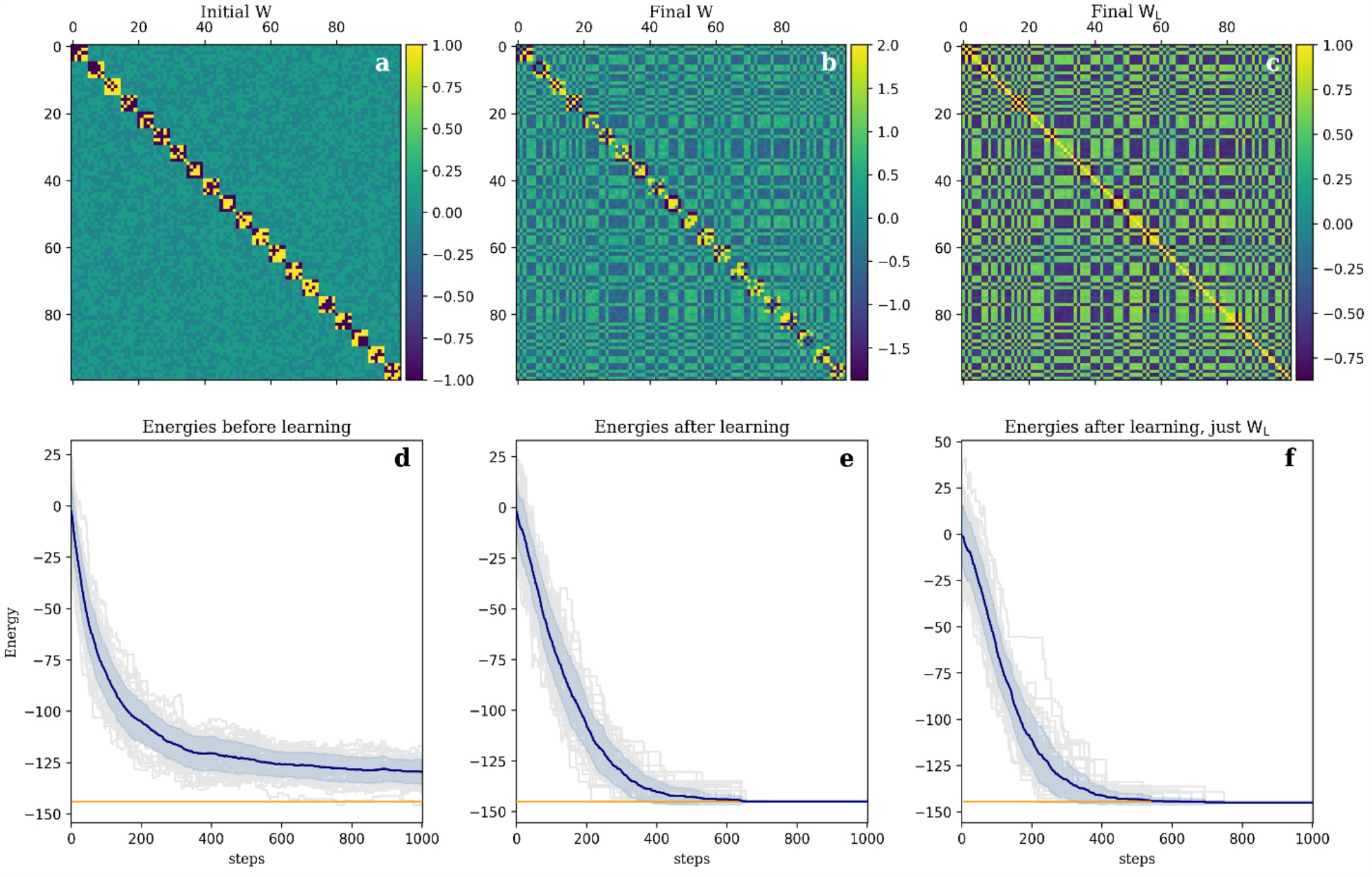
Dynamics of the SO model before and after learning. a) The original weight matrix **W**, b) the weight matrix W after learning, c) the learned part W_5_ of the weight matrix W. (d) Before learning; this is equivalent to the dynamics of a regular Hopfield network, where the system will converge to various minima according to the initial state. (e) The same system after the completion of Hebbian learning. We can see that for all initial random states the system converges to a lower energy state than it did before learning (indicated by the orange line). (f) The same system after the completion of Hebbian learning, but with state updates only driven by **W**_5_. On all plots d-f, the energy is computed using the original weight matrix **W** to see how the modified networks (b and c) perform on the original problem space (a), the state at each step is updated asynchronously, and the plots show the energy changes following 50 initial random states.

## 5.2 Stochastic Self-Optimization model -sinusoidal pulse train reset function *R*_1_(*t*)

In this section we present the results of resetting the system based on a sinusoidal pulse train for varying values of the pulse’s width and amplitude. Similar simulations were performed for a rectangular pulse train and the results are presented in Appendix A. In all simulations we examine a system with *N* = 100 nodes, a period *T*_0_ = 10*N*, 1000 pulses and the same weight matrix and learning rate as in Sec. 5.1.

Figure 7 shows the distributions of energy of the system that went through resets based on a sinusoidal pulse train reset function that has 1000 pulses, each pulse of width 40 and amplitude 3. Meaning that for this specific reset function at each pulse a maximum of *n* = 3 discrete states *s* of the system’s state ***S*** were reset at random to either 1 or -1 for a period of 40 steps.

**Figure 7.**
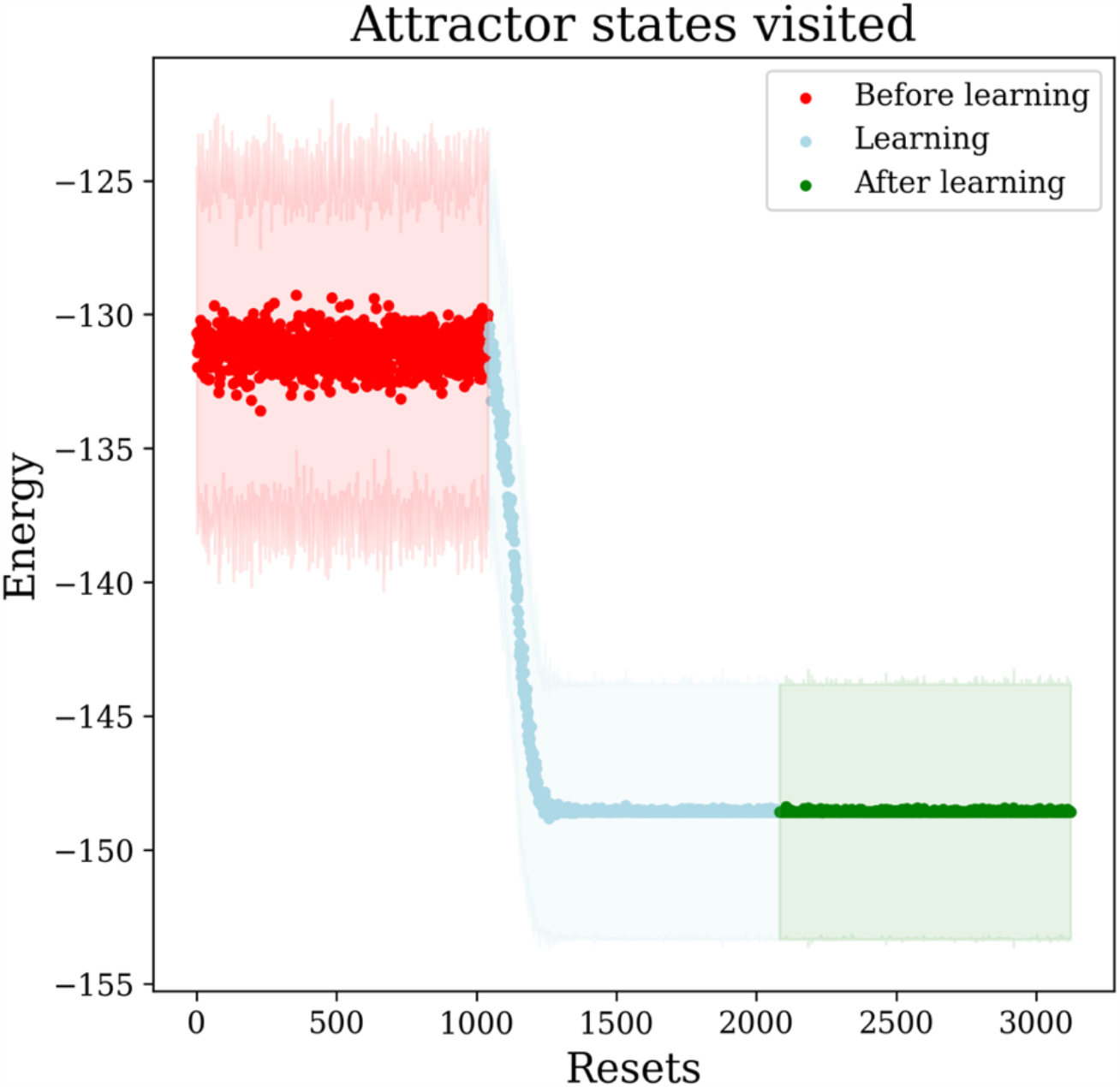
Energy of attractors visited when resets are a sinusoidal pulse train, where each pulse of width 40 and amplitude 3. The points represent the mean energy for 100 random runs at the end of convergence for a set without learning (resets 1–1000, red), during learning (1001–2000, blue), and after learning (2001-3000, green). The shaded areas in corresponding colors denote the standard deviation of the distribution.

We can see that for this configuration of the pulses, roughly after 300 pulses the system on average converges to an energy *E*_mean_∼ − 148. This final mean energy is similar to the case of instantaneous resets in the original SO model. However, since each pulse has a width of 40 steps, it means that it took the system (10*N* + 40)300 = 312,000 time steps to reach convergence. This is relatively faster than the original SO model, that converged after ∼10*N* × 520 = 520,000 time steps. This is further demonstrated in Fig. 8 that shows the converged energy during learning for various values of pulse width and amplitude after an increasing number of pulses.

**Figure 8.**
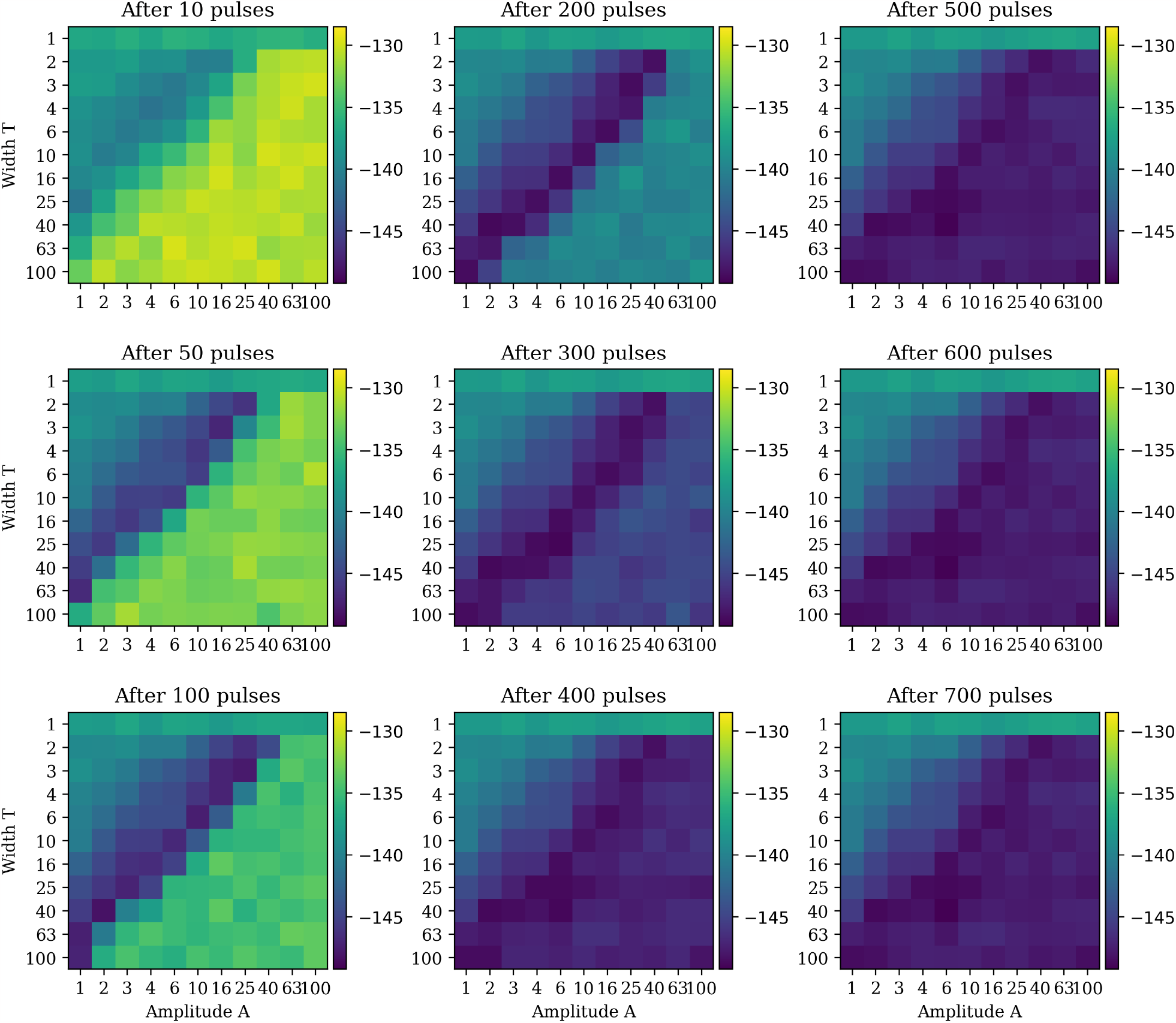
Resets with a sinusoidal pulse train. Each plot shows the mean energy of attractor states visited (100 random runs) during the learning phase (blue range in Fig. 7) for varying pulse width T and amplitude A (11 values for each ranging from 1 to 100 in a logarithmic scale) after different amount of pulses.

Figure 8 shows the mean energy of attractor states visited for the learning phase (the blue range in Fig. 7), for the sinusoidal pulse train of varying pulse width and amplitude, after different amounts of pulses (i.e. resets). For reference, the values of the learning phase in Fig. 7, where the pulse train has a width of *T* = 40 and amplitude *A* = 3, appear in row *T* = 40 and column *A* = 3 in each of the subplots of Fig. 8. We can see that after 100 pulses the system converges to an attractor that is around *E*_mean_∼ − 142, and after 200 pulses it is reaching *E*_mean_∼ − 148.

**Figure 9.**
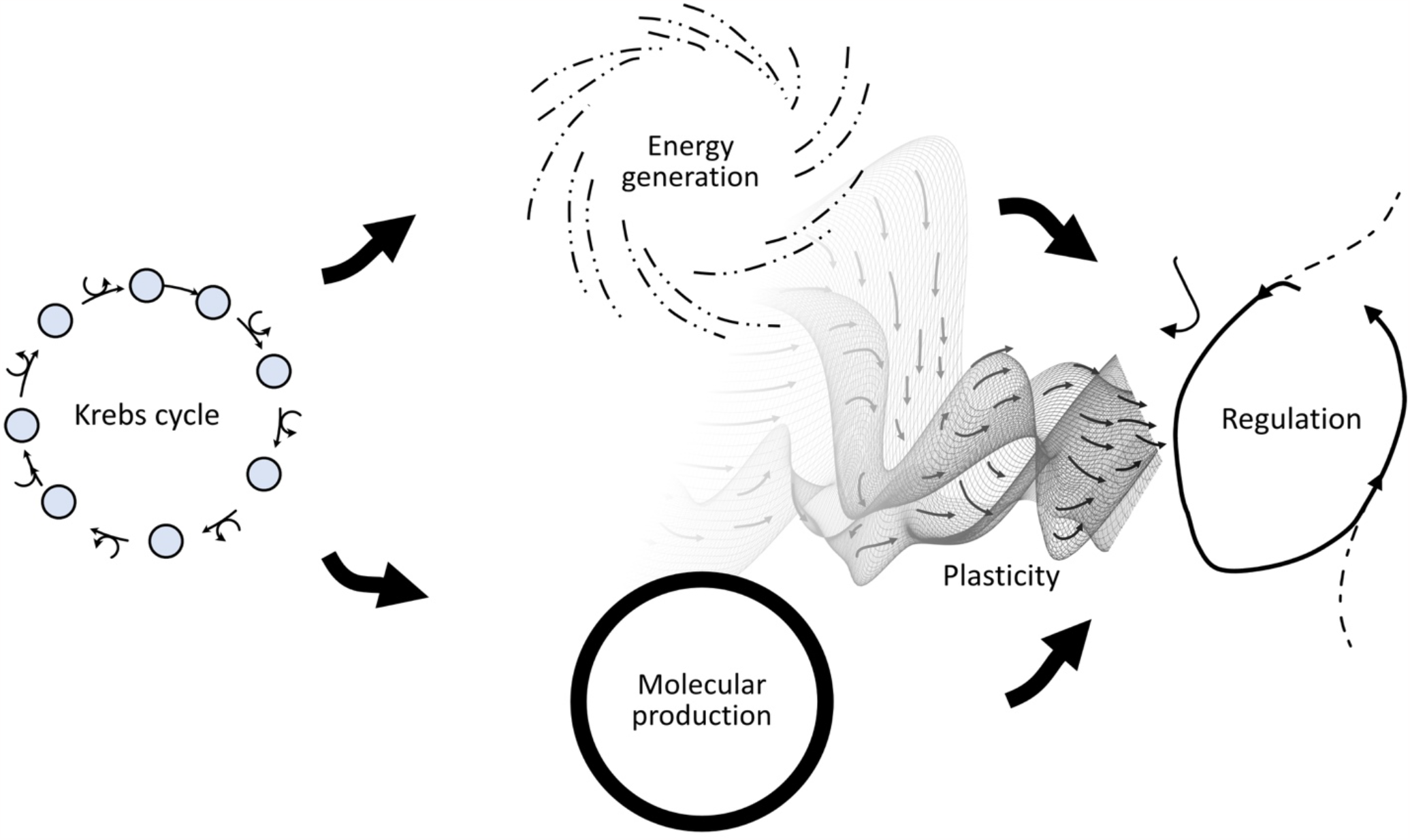
A working hypothesis about the paradoxical organization of the Krebs cycle. On the left, is the Krebs cycle, which embodies a tension between two mutually exclusive modes of operation: energy generation (middle, top) and molecular production (middle, bottom). The SO model suggests that switching back and forth between these two modes could spontaneously give rise to a process of coordination capable of generalized constraint satisfaction, as long as there is plasticity allowing for the accumulation of associative memory.

We can note several things. First, we should be able to recover the instantaneous reset by looking at a pulse of width *T* = 1 and amplitude *A* = 100, since that would be the analogous of resetting the entire system in one time step. And, indeed, if we look at the Fig. 10 in Appendix A showing the rectangular pulse train, we can see that the system reaches the mean energy *E*_mean_∼ − 148 after about 500 pulses (compare to Fig. 5, blue range). It should be noted that in Fig. 8, the row for amplitude of *A* = 1 is an artifact of trying to fit a sine peak into a discrete time step of width 1, this is avoided in the case of rectangular pulse train.

**Figure 10.**
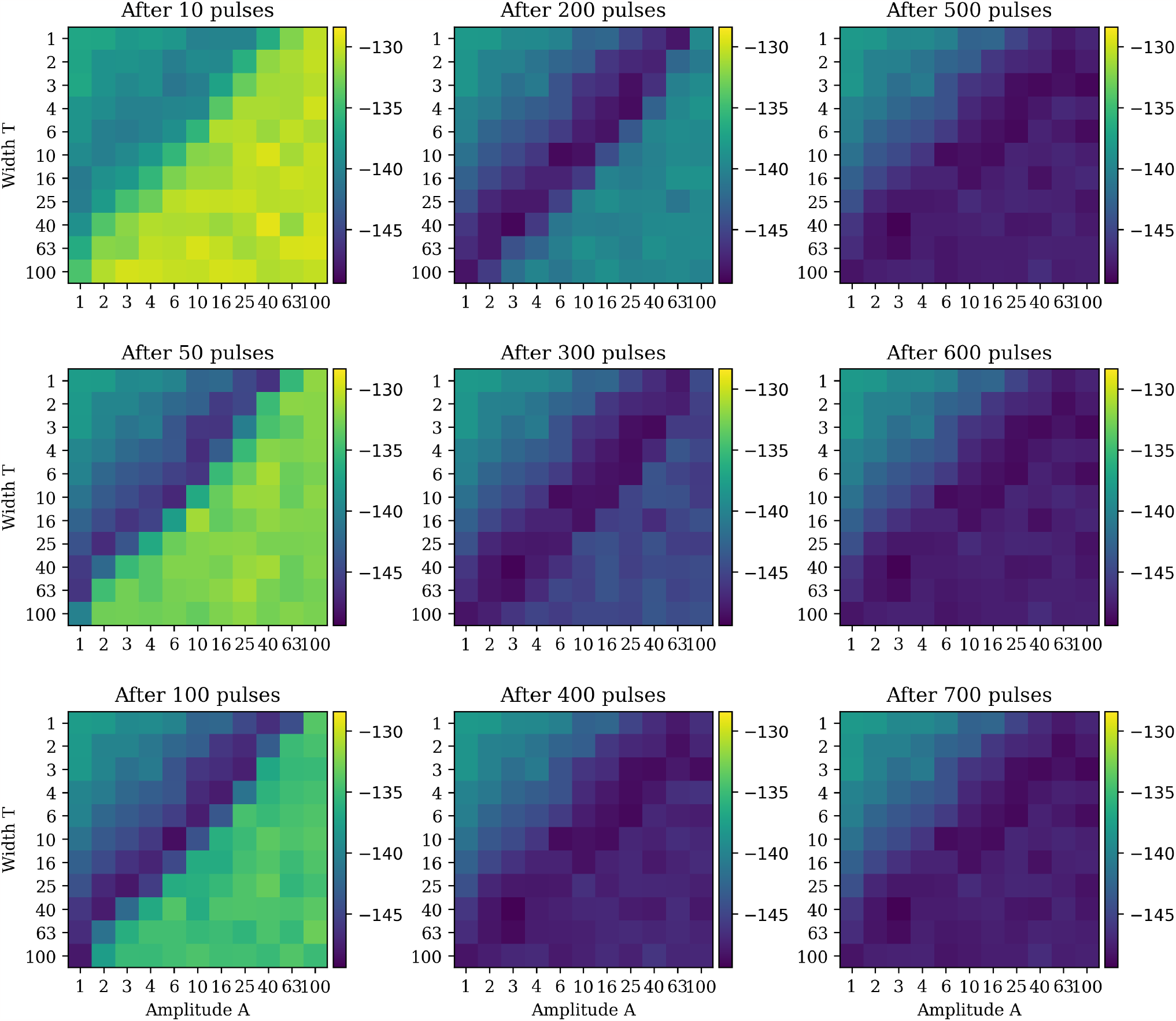
Resets with a rectangular pulse train. Each plot shows the mean energy of attractor states visited (100 random runs) during the learning phase (blue range in Fig. 5) for varying pulse width T and amplitude A (11 values for each ranging from 1 to 100 in a logarithmic scale) after different amount of pulses. We can see that the analogous state to the original instantaneous reset model, that is a pulse of width T = 1 and amplitude A = 100, indeed reaches the mean energy E_678W_∼ − 148 after 500 pulses.

Second, after 400 pulses, there is full convergence for all the different sizes of the pulses. Third, the straight-line minimum in each of the subplots in Fig. 8 indicates that there is an inverse relationship between best width and best amplitude, i.e. area is the deciding factor. We can see that when the area of the pulse is roughly of the size of the system (in this case *N* = 100), the system converges to lower energy values already after 200 pulses. If the area of the pulse is lower than the size of the system, the system converges to a local minimum instead. If the area of the pulse is larger than the system size, it will take the system longer to reach the global minimum. Therefore, width times amplitude on the order of *N* appears to be the most efficient at reaching the lowest converged energy in the fewest resets. To make this clearer we show in Fig. 11 in Appendix B the time dependence for parameter combinations on the diagonal of subplots in Fig. 8 for all time steps. For pulse areas larger than the system size the lowest converged energy is not always reached; the reason for this is not clear and is subject to future investigation.

**Figure 11.**
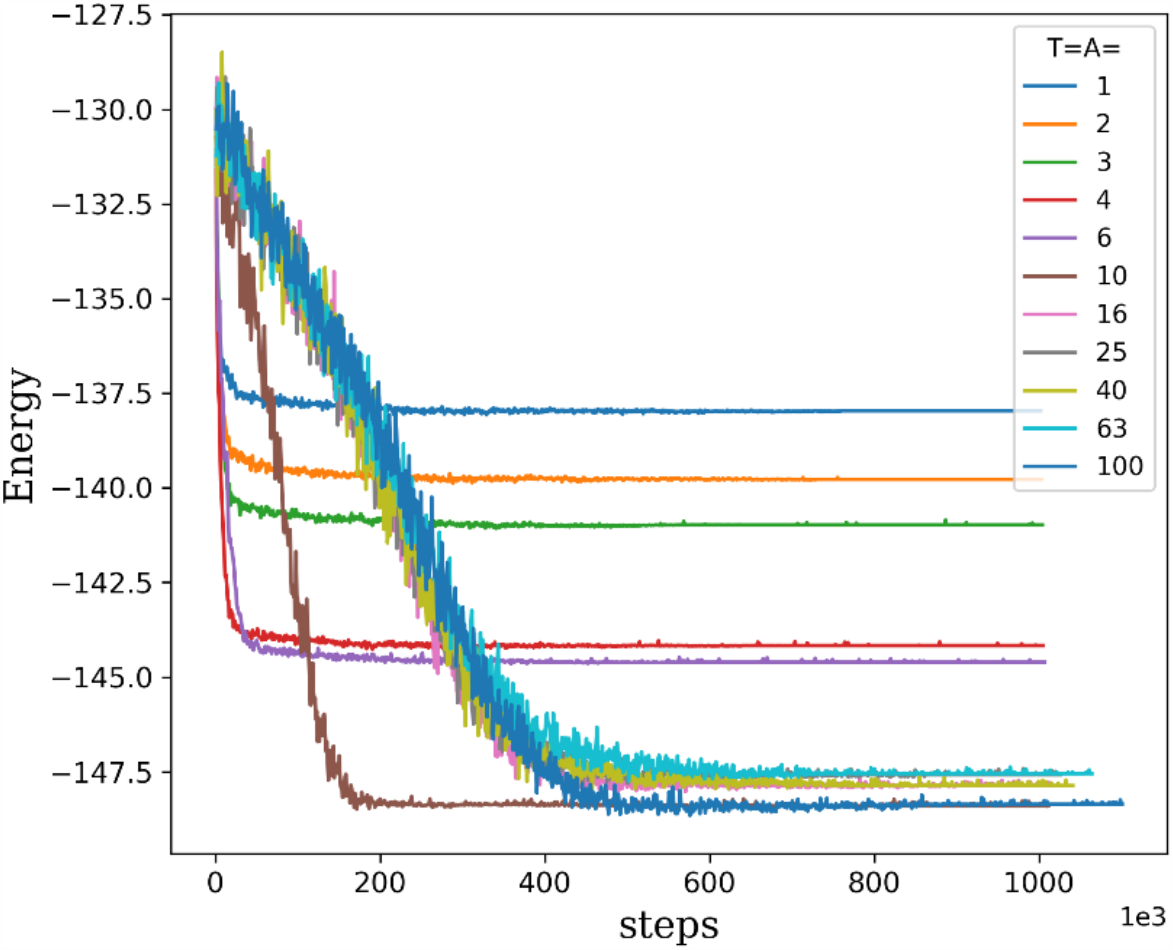
Resets with a sinusoidal pulse train: Energy of attractors visited as a function of reset steps for pulses of increasing width and amplitude such that T = A (values given in the legend). This corresponds to the diagonal of the subplots of Fig. 8.

## 5.3 Summary of modeling results

This probabilistic SO model, like the original SO model, acquires an improved capacity for constraint satisfaction. Overall, the results have demonstrated the following:

1. As expected, when the Hopfield network is repeatedly pushed far from of its equilibrium state, Hebbian learning will induce an associative memory of state regularities that is the basis of generalized constraint satisfaction.
2. As expected, this acquired memory can stand in for the optimized Hopfield network as a whole and does equally well in constraint satisfaction of the original problem space.
3. These results are quite general; in addition to the different implementations explored in the literature, here we showed that they do not depend on the precise way in which the system is pushed away from equilibrium; partial resets work too.
4. The results indicate that a probabilistic reset of system states could be preferable in terms of the temporal efficiency of self-optimization, compared to a complete instantaneous reset of all states.

In summary, the SO model illustrates how a system that is digging deeper into the trajectories through which it passes, while being driven through a sequence of configurations, can give rise to spontaneous re-structuring such that it acquires a more generalized constraint satisfaction capacity. This capacity can be an emergent property based on changes at the system level, but it can also be based on a dedicated a sub-structure within the system. Moreover, the fluctuations that drive the system do not need to be extreme to achieve this result; a repeated partial loss of the closed-state equilibrium state is sufficient and perhaps even more efficient in terms of the time required to adapt.

## 6 Discussion

The connection of the SO model with the enactive approach to autopoiesis with adaptivity under precarious conditions opens a rich source for further development. In order to make these lines of work relevant for mainstream biology, future work should focus on making the model less abstract and link it more closely with work on actual living systems. Here we will sketch a few avenues that could become topics for future work.

### 6.1 Scaling

For illustrative purposes we have worked with a relatively small Hopfield network of a hundred nodes. A concern might be that indeed most research on the self-optimization model has focused on network sizes on the order of a hundred nodes, so that it is unclear whether the process scales over several orders of magnitude. We recently put this concern to rest by demonstrating that the self-optimization model can be scaled up to ten thousand nodes with comparable results (N. Weber, Koch, & Froese, 2022). This scalability opens an intriguing perspective on growth.

It is well known that body growth leads to differential scaling of body parts. A classic example is allometric scaling of volume-based properties compared to length-based properties, the latter of which only scale with body mass to the power of 1/3. This implies, for example, that the length of neurons cannot keep up with brain volume, such that growth naturally results in sparser brain connectivity. Allometry can therefore provide a geometric account of the well-known fact that modularity increases with brain size, which in turn offers cognitive advantages (Changeux, Goulas, & Hilgetag, 2021).

Future work could implement a spatially embedded version of the SO model by taking inspiration from this size-dependent modularity of biological networks. When there is little sparsity or modularity to exploit, the best that the SO model can do is *selection* and *recollection*: it starts to preferentially converge on the deepest equilibria from among those that it has encountered in the past. When modularity is prominent, it can do even better by *generalization*: it will associate features from past equilibria and thereby move toward areas of state space that are otherwise practically inaccessible and where new, even deeper equilibria can be found. In other words, we expect that as a spatially embedded SO model would grow from a fully interconnected network to a sparser network, its capacities spontaneously would shift from mere basic coordination to more complex cognitive capacities – without adding extra requirements on the mechanisms already involved.

### 6.2 Regulation

The structural changes that the system acquires in its weight matrix due to Hebbian learning can, once sufficient changes have accumulated, by themselves *stand in for* the whole optimized system (i.e., original weights combined with the acquired weights) to solve the original constraint satisfaction problem. This is an important capacity because it confirms that the basic ingredients of the SO model result in a system that satisfies a key theorem of cybernetics and control theory – the good regulator theorem – which states that “every good regulator of a system must be a model of that system” (Conant & Ashby, 1970). This finding is also consistent with the original accounts of this optimization process that emphasized its capacity for the “self-modeling” of its own attractor landscape (Watson et al., 2009; Watson, Buckley, et al., 2011).

This dual possibility of solving the original constraint problem by using either the whole optimized Hopfield network, or just the subset of changes that resulted from the self-optimization process, is intriguing. It is reminiscent of a controversial claim about how to interpret generative models in the context of the free energy principle framework, namely that “an agent does not have a model of its world—it *is* a model” (Friston, 2013, p. 213). This validity of this ontological claim has been much debated, and a more cautious, instrumentalist interpretation of this notion of model can be articulated (e.g., van Es, 2021). Still, even with an instrumentalist stance we may wonder whether to attribute that status to the whole (the system is a model) or some of its parts (the system has or contains a model). What is intriguing about the self-optimization process is that it does not matter: both ways of carving up the system/model relationship work equally well.

Another point of concern is the representational status of the concept of a model. In cognitive science the default interpretation of a system’s model is in standard representational terms, i.e., that the model is representing something else, which is a function at which it can succeed or fail, or at least be better or worse at, etc. In other words, such an interpretation of a model as standing for a phenomenon brings normative questions into play having to do with evaluation of whether the model is doing a good job at representing the phenomenon which it is modeling. However, in general, naturalizing this representational content, such that its origins can be explained, is still a fundamental problem for cognitive science, and one that remains without a widely accepted solution (Hutto & Myin, 2013). To avoid any unintended representational interpretations, we will therefore continue to refer to this process as self-optimization instead of self-modeling.

Fortunately, there is no need to drop the concept of model altogether because there are simpler interpretations. In the context of the free energy principle, for example, it has been argued that the generative model “uses exploitable structural similarities encoded in the internal states of the organism”, yet without being a structural representation as such (Ramstead, Kirchhoff, & Friston, 2019, p. 236). Similarly, according to the enactive approach, the concept of a model does not entail a commitment to a representational interpretation: a model that works by being a structure that stands in for another structure has no need for representational content. This applies for instance to many human tools that serve as models (Noë, 2015), and also to neural re-use and other properties of the brain (Hutto & Myin, 2014). Thus, while we can cautiously talk about the development of a “model” during the self-optimization process, it is good practice to keep in mind that this is meant to be interpreted as something standing *in* for something else (structurally), rather than as something standing *for* something else (referentially).

This initial point of contact between the SO model and the free energy principle deserves to be investigated more systematically in future work. The self-optimization model may not be capable of all the functionality that a generative model of the free energy principle offers, but it also has less requirements to get off the ground. Importantly, it does not rely on any error signals to develop its internal modeling capacity.

### 6.3 Entropy

The self-optimization process, when applied to a system with exploitable regularities, reliably reorganizes such that it finds state configurations of increased coordination than it would otherwise encounter. Moreover, the process makes the system reach these configurations at a *faster* pace. The analogy between the Hopfield network and a thermodynamic system is admittedly rather lose; nevertheless, the Hopfield network’s tendency to re-organize its connectivity such that it satisfies its constraints more efficiently is reminiscent of the large-scale spatiotemporal reorganization associated with the law of maximum entropy production (LMEP) (Swenson, 1997), also known as the fourth law of thermodynamics (Morel & Fleck, 2006; Swenson, 2009).

Consider, for example, the emergence of Rayleigh-Bénard convection rolls in a pan of oil that have the effect of organizing fluid flows such that heat transfer from the fire below to the air above is increased. Classic thermodynamics describes the constraints on a system’s direction of change, but it does not place constraints on how or at what rate that change is achieved. What the LMEP principle adds is that, in a multistable system with all else being equal, the system’s steady state with the highest rate of entropy production is favored. This claim has now been formally confirmed (Endres, 2017). The insights of LMEP have also been restated by “constructal theory,” which holds that “for a finite-size flow system to persist in time (to live) it must evolve such that it provides greater and greater access to the currents that flow through it” (Bejan & Lorente, 2010, p. 1335). Future work could more formally work out whether the SO model also falls into this class of systems, which would depend on finding a way to properly measure the entropy production associated with state changes in a Hopfield network.

However, from the perspective of the enactive conception of life, a complete reduction of the SO process to a purely physical phenomenon would be counterproductive for the characterization of specifically living systems. It would mean losing sight of what is distinctive of living compared to other material systems, which also has been an issue for the free energy principle (Kirchhoff & Froese, 2017). Accordingly, ecological psychology has emphasized a key difference between generic autocatakinetic (“ACK”) systems – systems that are characterized by self-production in addition to the MEP principle – and ACK systems that are additionally cognitive:

The hallmark of cognitive systems is that they can constitute their ACKs in ways that are arbitrary to and often antithetical to local force-fields or potentials. Their end-directed behavior is determined not by local potentials but by meaning. They use their on-board (stored) potential to disconnect from, or ignore, and move across local force-fields to search out, discover, and access new and discontinuously located potentials using the invariant (symmetry) properties of ambient energy flows that nomologically provide information about paths to distal ends or potentials. (Swenson, 2020, p. 106)

This description of a cognitive ACK system comes close to key ingredients of the self-optimization process, especially given that the system’s repeated transitions between configurations are achieved by a “reset” that temporarily makes state dynamics operate in a way that is “antithetical to local force-fields or potentials”. The SO model therefore provides a novel interpretation for why cognitive or biological ACK systems tend to be characterized by the kind of rhythms that are highlighted by Mahulikar and Herwig (2004): “This requirement to continuously reverse back to earlier state through non-linear processes is what leads to an irreversible loss. These cyclic processes involving ordered sub-systems increase the entropy of the surroundings at a faster rate.” (p. 219). What the SO model suggests is that periodic reversing of energy levels is also the motor that drives the system’s capacity to discover discontinuously located potentials. Future work could explore this new link between the enactive approach and ecological psychology in terms of the SO model and the cognitive ACK system.

## 7 Toward a workable model of biological regulation

Our hope is that by elaborating the enactive conception of life we have moved this framework in a direction where it can more productively connect with open questions in the biology of adaptive regulation. This elaboration consisted in two parts: (1) we advanced the enactive approach to autopoiesis by articulating the relationship between life’s precariousness and adaptivity, and (2) we introduced a way of formally modeling this relationship in terms of the SO model.

The starting point was that the primordial tension afflicting self-individuation – between a living system’s need for both openness and closure – is not a contingent fact about precariousness of life as we know it, i.e., which could be potentially avoided under other circumstances. On the contrary, as suggested by the SO model, the system’s sedimented itinerancy between these two conflicting requirements could be the very basis of adaptive regulation. The relationship between precariousness and adaptivity can therefore be considered as another key example of what Jonas ([1966] 2001) called life’s “needful freedom”. He exemplified this notion in terms of metabolism – the system *needs* a flow of matter and energy such that its organization can be *free* from any specific configuration of matter and energy. And he argued that this principle applies across all levels of organic organization. To take another example, an animal needs to move to feed such that it can be free to move.

Similarly, we suggest that the primordial tension of self-individuation is what enabled the first living systems to be capable of regulating that very same tension. Indeed, the emergence of this primordial capacity for self-regulation can be considered as defining moment at the origin of life itself – it is that which distinguishes living systems from other far-from-equilibrium self-producing systems. The evolution of higher levels of complexity did not necessarily reduce the tension of self-individuation; rather, new forms of life may have internalized the tension and thereby at the same time opened new possibilities of self-regulation. The minimal conditions for the SO model – closed dynamics, open dynamics, and plasticity – lead us to suspect that the SO process can be found in many places in biology exhibiting rhythms between opposite requirements.

The basic conditions were already recognized as prevalent by proponents of the LMEP: “ordered sub-systems are characterized by a low tendency towards dispersion relative to their surrounding disorder, through periodic coalescence and concentration of energy and matter for localized control of dispersion” (Mahulikar & Herwig, 2004, p. 214). They mention the “cyclic process of breathing and metabolism” as an illustrative example and highlight that such “cyclic processes oscillate the state of ordered sub-systems about the equilibrium (with disorder) state” (p. 219). The SO model can therefore be interpreted as an abstract model of the role of time-varying disordering in adaptivity. Moreover, in contrast to Mahulikar and Herwig’s denial that ordering through a cyclic process such as breathing and metabolism can be a manifestation of “purposeful activity” (p. 220), future work could even consider motivated activity precisely as another source of time-varying disordering enacted by living systems (Froese, 2023; Froese & Karelin, 2023).

This brings us back to the specific biological example that framed the start of this article, namely the metabolic tension inherent in the organization of the Krebs cycle – molecular production versus energy generation. As Lane (2022, pp. 19-20) puts it: “why use the same pathway to create and destroy, to burn and renew?” The SO model suggests a novel perspective: this cycle could be an internalized instantiation of the primordial tension of self-individuation between openness and closure, which was also grounded in competing needs of energy and matter but where that tension was largely a spatial one of inner versus outer. Hence, having these two kinds of internal needs linked into a *cycle* would guarantee that there are still two mutually exclusive modes of activity. The only missing ingredient for a SO process based on the Krebs cycle is plasticity: its itinerant path between the two modes would need to leave a trace such that its activity changes in a precedent-based manner. Interestingly, this requirement aligns with Lane’s rejection of a gene-centered view; for him it is the unceasing flow of energy and matter through a cell that “conjures the genes themselves into existence”, and shapes their activity of gene expression such that it is “*the movement that creates the form*” (2022, p. 6; emphasis added). Lane’s view and the SO model complement each other, as the latter has been linked to associative memory in the context gene regulatory networks (Watson, Buckley, Mills, & Davies, 2010; Watson et al., 2014).

These conceptual links between the SO model and fundamental questions in cell biology deserve to be more fully explored in future work. What is particularly noteworthy is that these links make it possible to sketch new theoretical perspectives on specific adaptive functions of otherwise puzzling features of cellular organization and dynamics. For instance, the requirement of the SO model to be rhythmically driven through a sequence of configurations could be developed into a novel perspective on the puzzle of why gene expression tends to occur in bursts, a phenomenon known as transcriptional bursting (Tunnacliffe & Chubb, 2020). This shows that the enactive conception of life in terms of self-individuation, now elaborated in terms of the process of self-optimization and implemented in a mathematical model, has the potential for connecting more productively with theoretical and experimental biology.

## Data availability

The code that was used for the implementation of the original SO model simulations is available in an online repository at: https://github.com/nata-web/SO-scaled-up. For the simulations of the stochastic SO model, i.e., resetting with sinusoidal or rectangular pulse train reset functions, Eqs. (5-7) were used instead of instantaneous resets.

## Declaration of competing interest

The authors declare that they have no known competing financial interests or personal relationships that could have appeared to influence the work reported in this paper.

## Funding

This work was supported by JST Grant Number JPMJPF2205. The funding source was not involved in the study design; in the collection, analysis and interpretation of data; in the writing of the report; nor in the decision to submit the article for publication.

## Author contributions

TF and TI initiated the development of the conceptual framework. NW and IS contributed to conceptualization. TF wrote the first draft of the manuscript. NW carried out the simulations and wrote the sections describing them. All authors reviewed, contributed to the article and approved the submitted version.

## Acknowledgements

TF thanks the organizers and audience of the *Workshop on Life Mind Continuity*, held on Nov. 25, 2022, at OIST in Okinawa, Japan, where an earlier version of the ideas related to the Krebs cycle were presented. TI is grateful to OIST’s Theoretical Sciences Visiting Program, which provided the context for this collaboration. All authors are grateful for the help and support provided by the Scientific Computing and Data Analysis section of the Research Support Division at OIST.

## Appendices

*Appendix A - Rectangular pulse train, R*_*rect*_(*t*)***W***

*Appendix B – Time dependence for parameter combinations*

## Notes

https://github.com/nata-web/SO-scaled-up

### Competing Interest Statement

The authors have declared no competing interest.

### Summary of Updates

This is a substantially revised version of the manuscript that clarifies the links between the enactive conception of life, the self-optimization model, and thermodynamics.

